# Persistent loss of biologically-rich tropical forests in the Indian Eastern Himalaya

**DOI:** 10.1101/827360

**Authors:** Chintan Sheth, Aparajita Datta, Devathi Parashuram

## Abstract

Deforestation is a major cause of biodiversity loss in Asia. Using fine-resolution satellite imagery we assessed the change in forest cover of a state-managed Reserved Forest located in India’s Eastern Himalaya biodiversity hot-spot. Thirty-two square kilometers of forest cover was lost from 2013 and 2017 with a 5% decline in total forest area over four years. Hornbills are a key functionally important species found in the area. We therefore assessed the habitat around 29 hornbill nest trees in this Reserved Forest and estimated that there was a loss of 35% of forest cover from 2011 to 2019. We identify illegal logging (despite a ban by the Supreme Court of India) as the main driver that is depleting forest cover within this important area. Our results highlight the ongoing threats to biologically-rich forests and the need for urgent measures to halt this loss. We suggest that this study has practical implications for the governance of non-PA state-managed forests in Arunachal Pradesh. The ongoing deforestation appears to be due to organized crime, institutional inadequacy from a combination of limited resources, bureaucratic apathy, and/or ambiguity in use and ownership of forest land compared to other community forests which appear to have robust governance systems.

## Introduction

Tropical forests are not only the most biodiverse terrestrial ecosystems on Earth (Gibson et al. 2011) but also amongst the most threatened. Globally, 2.3 million square kilometers of forest were lost from 2000 to 2012, with tropical forests undergoing the highest losses (Hansen et al. 2013). Deforestation is one of the major causes of biodiversity loss across the world (Gibbs et al. 2010; Curtis et al. 2018).

India’s state of forest is assessed biennially by the Government’s Forest Survey of India (FSI) and according to FSI, India has lost 80% of its native forest cover and forests continue to be lost at the rate of 1.5 to 2.7% per year. However, this does not provide an accurate estimate of the true extent of native forests and deforestation rates as these data combine native forests, secondary regrowth, plantations and cropland and do not validate classifications with ground-truthing (Puryavud et al. 2010a, b). Puryavud et al. (2010b) highlighted the cryptic destruction of India’s native forests as a challenge to understanding the trends in the state of India’s forests.

Global Forest Watch (GFW) data show that India lost about 15,400 km^2^ of forest (>30% canopy cover) between 2001 and 2017 amounting to 172 mega tonnes of CO_2_ emissions (Hansen et al. 2013; Global Forest Watch 2019). North-east India, which encompasses two global biodiversity hotspots – Indo-Burma and the Himalaya (Mittermeir et al. 2005) – appears to be severely affected by deforestation (Pandit et al. 2007). The GFW assessment estimated 11,400 km^2^ of forest loss from north-east India in the same period (Global Forest Watch 2019).

Arunachal Pradesh in north-east India is the richest terrestrial biodiversity region in India (Mishra & Datta 2007) with nearly 6000 flowering plants and half of the bird species known from India (Praveen et al. 2016, 2019). Recent research has led to the discovery of new records, range extensions and new species of plants and animals from the state (Gajurel et al. 2001; Ahti et al. 2002; Ahmad et al. 2004; Sinha et al. 2005; Athreya 2006; Tamang et al. 2008; Sondhi & Ohler 2011; Zanan & Nadaf 2012; Dalvi 2013; Roy 2013; Hareesh et al. 2016; Siliwal et al. 2017; Captain et al. 2019).

Forest cover in Arunachal Pradesh has been declining in the last decade although forests still cover 79% of the total land area (Global Forest Watch 2019; Supplementary Table 1 & Supplementary Figure 1). FSI reports estimate that 486 km^2^ of forest was lost from 2003 to 2017 in Arunachal Pradesh (FSI 2003, 2005, 2009, 2011, 2013, 2017). However, GFW data, shows that 2000 km^2^ of forest was lost between 2001 and 2018, comparable to a 3.2% decrease in forest cover since 2000 (Global Forest Watch 2019, Supplementary Table 2).

In terms of their legal status, 11.37 % percent (9528 km^2^) of the geographical area of Arunachal Pradesh is under the Protected Area (PA) network (Wildlife Sanctuaries and National Parks, some of which also encompass Tiger Reserves). The PAs are generally better protected than Unclassed State Forests (USF; 37% of area; 30,965 km^2^) and Reserved Forests (RF; 11.61% of area; 9722.69 km^2^) attributable to stronger implementation of the country’s forest and wildlife laws. USF areas are in practice used and/or owned by the community (*de facto* rights), although recorded as being under the Forest Department. The RFs despite being legally under the control of the state Forest Department are often subject to various anthropogenic pressures such as agricultural expansion, conversion to plantations and/or logging (Naniwadekar et al. 2015a).

With 80% of the population practicing subsistence farming in the hilly terrain, people were primarily dependent on shifting cultivation which is mainly carried out in the USF or community forests. Shifting cultivation was estimated to cover 2040 km^2^ in 2008-09 (Wasteland Atlas 2011) but is now in decline among many communities (Teegalapalli & Datta 2016). Although shifting cultivation is usually cited as the main driver of forest loss in the state, there are several drivers of forest loss such as: agricultural expansion, growth of plantation crops such as oil palm, rubber, tea, opium, illegal logging and road expansion (Srinivasan 2014; Velho et al. 2016; Khandekar 2019). With an increasing population, need for agricultural land and development, and lack of land demarcation and cadastral surveys, there is logging (mostly in Reserved Forests) for agriculture expansion and plantations along with illegal logging in Arunachal Pradesh (Naniwadekar et al. 2015a; Velho et al. 2016; Rina 2017, 2019; Mamai 2018; Khandekar 2019).

### Ethno-civil conflict and illegal logging

The main sources of revenue for Arunachal Pradesh were forest-based industries till 1996, after which the Supreme Court banned logging. Despite the ban, illegal clearing driven by ethno-civil conflict in Sonitpur district in neighbouring Assam resulted in the disappearance of several Reserved Forests that bordered Nameri Tiger Reserve in Assam in the last two decades (Srivastava et al. 2002; Kushwaha & Hazarika 2004; Mazoomdar 2011; Velho et al. 2014; Srinivasan 2018). Srivastava et al. (2002) estimated that 232 km^2^ of forests was cleared in Sonitpur District between 1994 and 2001 with the overall loss rate of 28.65%, possibly the highest deforestation rate in the country. Kushwaha and Hazarika (2004) estimated 344 km^2^ forest loss between 1994 and 2002 in the Kameng and Sonitpur Elephant Reserves, while Velho et al. (2014) reported continuing forest loss in the same region around the southern boundaries of both Pakke and Nameri Tiger Reserves. Between 2001 and 2018, 170 km^2^ of forest was lost from Sonitpur district (Global Forest Watch 2019). Forest loss over twenty-five years has resulted in substantial habitat loss for wildlife that include tigers, elephants and large birds such as hornbills.

After the 1996 ban, selective logging has re-started in some forest divisions in Arunachal Pradesh since 2008-2009. However, apart from these state-controlled and permitted logging activities, ground observations and local media reports indicate that illegal logging is becoming a major driver of deforestation in Reserved Forests (Rina 2017; Anonymous 2019) and other areas in Arunachal Pradesh (Mamai 2018; Anonymous 2019).

The Pakke Tiger Reserve and its surrounding Reserved Forest areas are among the few remaining areas of low-elevation forest and is among the best areas for hornbills in South Asia (Datta 1998, 2001; Datta & Rawat 2003, 2004; Dasgupta & Hilaluddin 2012; Datta et al. 2012; Datta & Naniwadekar 2015) due to protection measures by forest authorities (Velho et al. 2011) and control of hunting by local people. Hornbills are an ecologically important functional group that act as effective seed dispersers (Datta 2001, Naniwadekar et al. 2015, 2019a, b). The main nesting habitat for the Great hornbill *Buceros bicornis*, Wreathed hornbill *Rhyticeros undulatus* and Oriental Pied hornbill *Anthracoceros albirostris* lies along the low-elevation areas encompassing the Pakke Tiger Reserve and surrounding Reserved Forsets (Datta & Rawat 2004) where illegal logging occurs. In 2012, the Hornbill Nest Adoption Programme (HNAP) was initiated to protect hornbill nest trees and nesting habitat in the Papum Reserved Forest (Fig. 1) outside Pakke Tiger Reserve in a partnership with local communities and the state Forest Department (Datta et al. 2012; Rane & Datta 2015). Since the programme began, it has resulted in increased local awareness about hornbills and nest trees of three hornbill species have been protected with successful breeding and chick production. However, ground observations indicate increasing levels of illegal tree felling from 2016, with the use of mechanized chainsaws, hired labour from outside and the transport of timber outside the state.

**Fig. 1.**
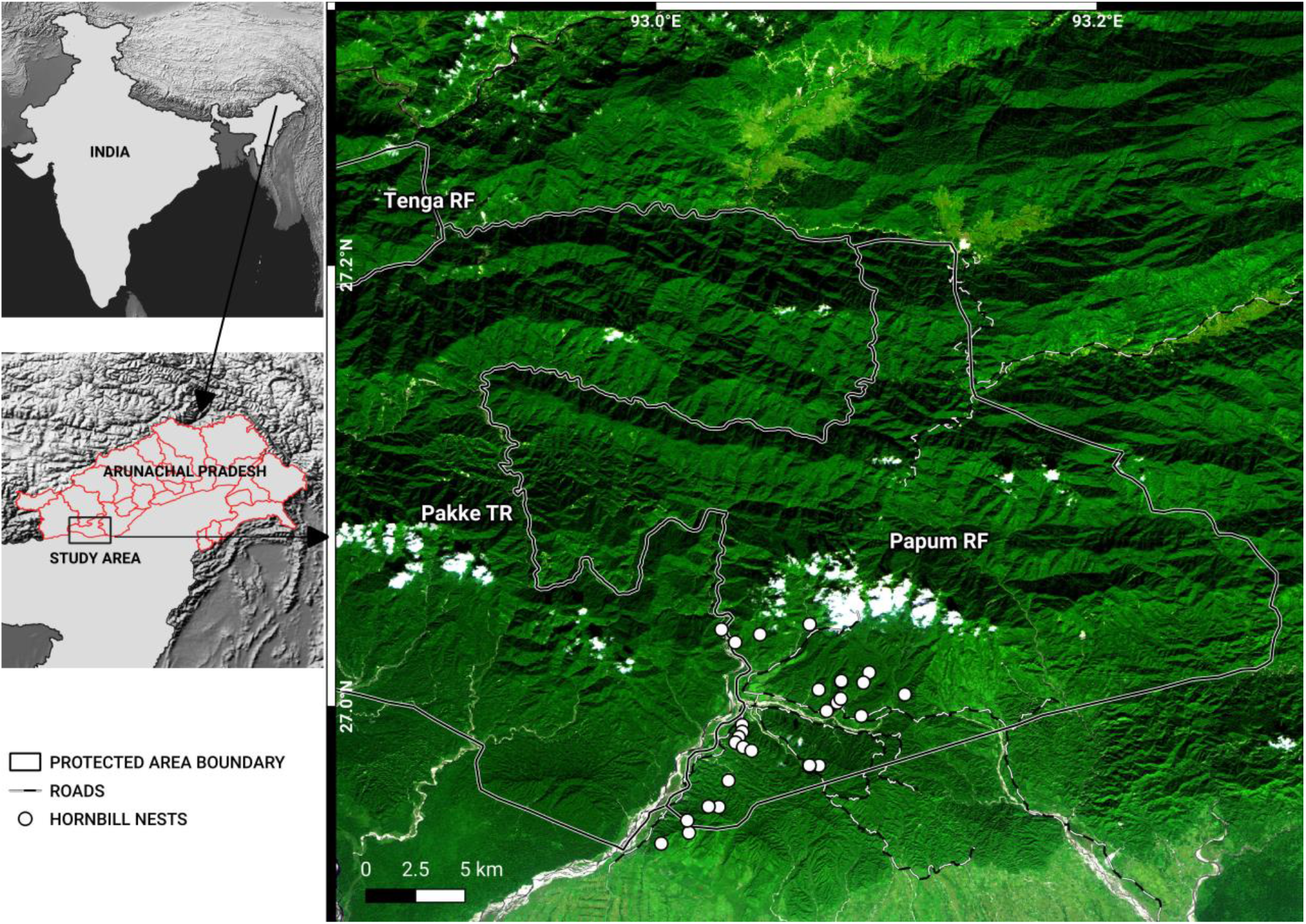
A November 2018, false-colour composite image (RapidEye bands 3, 4, 1) of the study area, showing Pakke TR, Tenga RF and Papum RF. The administrative boundaries of Arunachal Pradesh are marked in the map of India. The border between Assam and Arunachal is also the lower boundaries of Pakke TR and Papum RF. Shades of dark green indicate forests in different elevations. Lighter green shades are cropland, secondary vegetation, flooded grasslands and bamboo. Whites are indicative of clouds, river beds and landslides. Blue depicts water. Notice the density of roads in the southwest of Papum RF.

In this study, we aimed to 1) assess the extent of forest loss in the Papum Reserved Forest which adjoins the Pakke Tiger Reserve. Given that hornbills are a key faunal group that is functionally important and their main nesting habitat lies in the foothill areas which are affected by illegal logging, we also aimed to assess the loss of hornbill nesting habitat. Our specific purpose is to 1) estimate forest loss in the Papum RF using satellite data at a fine-scale resolution (3, 5 m) from 2013 to 2017 and 2) to determine forest loss within 1 km of hornbill nest trees at a fine-scale.

## METHODS

### Study area

Papum RF covers an area of 1064 km^2^ and adjoins Pakke Wildlife Sanctuary and Tiger Reserve (henceforth, TR) (Fig. 1; 861.95 km^2^, 92.5932° - 93.1006°N; 26.9351° – 27.2283°E). The area is part of the Eastern Himalaya Biodiversity Hotspot. Papum RF receives an average total annual rainfall of 2500 mm. Mean (± standard deviation) maximum temperature is 29.3°C (± 4.2) and the minimum temperature is 18.3°C (± 4.7). The vegetation is classified as the Assam Valley tropical semi-evergreen forest (Champion & Seth 1968). Papum RF has a similar floral and faunal composition to the adjoining Pakke Tiger Reserve.

The Papum RF was constituted as a Reserved Forests where all extractive activities are prohibited unless legally permitted (Indian Forest Act 1927). Part of Papum RF (346.25 km^2^) is included in the buffer area of Pakke TR as per the National Tiger Conservation Authority (NTCA 2012), India. Of this 318.25 km^2^ is forested zone, while 28 km^2^ is demarcated as multiple use area (NTCA 2012). Within Papum RF, there are 19 small towns/villages and settlements with a population of 3789 (2011 Census of India). Towards the south and east, Papum RF is bordered by Assam and Papumpare district respectively. To the west, lies the Pakke River and Pakke TR; and to the north are the community forests of Pakke Kessang. Nameri Tiger Reserve in neighbouring Assam state is contiguous with PTR in the south.

Although the total area is 1064 km^2^, for this study, we marked out an area of 737 km^2^ for classifying the forest and analysis of change in forest cover (Fig. 1). We restrict our analyses to 70% of the total area for two reasons: 1) the geographical focus of the HNAP program is within this area, 2) the boundary of entire Papum RF is uncertain and 3) the region of our analysis also forms part of the buffer area of neighboring Pakke Tiger Reserve. A digitized boundary of Papum RF (737 km^2^, including a 500-m buffer; 92.9209° - 93.2826°N; 26.9446° - 27.2116°E) was used for the analyses.

### RapidEye and PlanetScope satellite data processing and image classification

To conduct a supervised image classification, all satellite images were pre-processed by PlanetLabs before analysis. For example: ortho-rectified radiance/reflectance data of the RapidEye (5 m spatial resolution in 5 spectral bands) and PlanetScope (3 m spatial resolution in 4 spectral bands) constellations were obtained to ensure a complete cloud-free coverage of the Papum RF region (for a list of images analysed refer to Supplementary Table 3; refer Planet Labs Inc. 2019 for dataset descriptions and spectral bands). We used fine-scale satellite images for land-cover classification as this resolution can robustly resolve forest loss and other ecological phenomena below the 30-m scale (Hansen et al. 2013, Milodowski et al. 2017). Ortho-rectification (a process of image correction to account for irregular topography) is applied to ensure the same geographical region is analyzed year-to-year within a region of interest (ROI) (Tucker et al. 2004). Scenes were chosen if they were entirely cloud-free and taken by the same satellite on the same day, thereby preventing complications of image stitching and loss of information due to cloud cover. Datasets from both satellite constellations were combined to include the oldest possible year of fine-scale data (2011), and whenever RapidEye data was unavailable for analyses (example data after 2016).

Each satellite scene (or partial scene) was independently classified as forest, non-forest and logged-forest using the randomForest library 4.6-14 (Liaw & Wiener, 2002) in the R software for statistical computing (R version 3.3., R Core Development Team 2016). Ground-control polygons (GCPs) were identified within the three land-cover classes by using a combination of field sampling (using a global positioning unit) and Google Earth imagery. Forest regions comprised GCPs of closed canopy forests with little or no detectable anthropogenic disturbance. Non-forest regions comprised water bodies, grasslands, permanent settlements, sand bars and landslides. Logged-forest GCPs were defined using ground reports of active/past logging, studying satellite images at GFW deforestation hot-spots, and for roads, new clearings, plantations and fire scars. Logged-forest GCPs generally comprise areas previously under forest but currently with higher albedo than forest. The shape of the clearings is often geometrical and close to older forest clearings. Roads are linear in shape with the lower slope scarred with discarded debris. The training datasets of the above three classes consisted of at least 40 GCP’s and ~29 million pixels, per year.

Land-cover classification of the entire Papum RF using fine-scale data was only possible for the years (2013, 2014, 2017), where these scenes fulfilled the above coverage criteria. However, the forest loss analysis around the hornbill nest trees utilized images from 2011 – 2019.

### Land cover change around hornbill nest trees

The HNAP is confined to the lower and south-western parts of Papum RF (Fig. 1) that fall within Seijosa circle – from Darlong up to Jolly/Lanka in the north and towards the Mabuso 2/Margasso settlements to the east, within Pakke-Kessang district.

To investigate if the habitat around 29 protected hornbill nest trees were affected by forest loss, scenes that covered >90% of the hornbill nest sites were chosen. Cloud-free, single day scenes were available and could be analysed from 2011 to 2019. This allowed us to make comprehensive fine-scale forest loss estimations for 9 years. Cloud-free satellite images for all years were from November-December, except for 2018 and 2019 which were from April-May (dry season). During the dry season, secondary vegetation in clear felled areas is visibly dissimilar from primary forest. While we do not test for this difference, we think the visible difference may be attributed to the drying and browning of vegetation in the summer season when soil moisture and rainfall are low. Secondary vegetation in winter months (post-monsoon October - February) are visibly greener as the soil moisture is still high. An identical approach (to that used for classifying forest loss in Papum RF) was implemented to classify the area around 29 hornbill nests. A 1-km buffer was created and the satellite scenes were clipped to the buffered extent (48 km2). Three land-cover classes were defined (see above) comprising 20 GCPs and ~ 2 million pixels (RapidEye data) or ~ 5 million pixels (PlanetScope data, refer to Supplementary Table 3).

The spatial accuracy of the land-cover classification was assessed by manual checking of the scenes combined with a stratified random sampling method (Olofsson et al. 2014). A random sample of every land-cover class in each training dataset was used to test the accuracy of the classified image providing a bias corrected estimate of land-cover area in each class. The associated standard errors, prediction accuracy and rates of commission and omission errors were estimated as recommended by Olofsson et al. (2014). The prediction accuracy and standard error of the classification (for three years’ of RapidEye data) is 98.4 ± 3.0%. For forest loss estimates around hornbill nest sites, the prediction accuracy is 96.4 ± 7.5%. Accuracy statistics and confusion matrices for both Papum RF and the nest-sites are tabulated in the supplementary material (Supplementary Table 4 and Table 5).

Our analyses combined results from both satellite datasets as we found the prediction accuracies to be comparable. Image classification prediction accuracies were estimated using the methods recommended in Olofsson et al. (2014). A two-sample randomization test was performed on the distribution of all possible differences between accuracies of the observed years and then compared to the observed difference between the mean accuracies of the respective datasets (observed difference in mean accuracy = 0.03351667, *p*-value = 0.2457542; Manly 1991).

The annual rate of forest area loss was calculated on the classified land-cover images using a modified compound interest-rate formula from Puyravaud (2003) for its mathematical clarity and biological relevance:

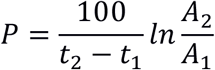

where A_1_ and A_2_ is the forest area in time periods t_1_ and t_2_, respectively. P is the annual percentage of area lost.

## Results

### Forest loss in Papum RF: 2013 – 2017

There was very high forest loss in Papum RF as determined from analysis at a fine-scale resolution. Table 1 shows the loss of forest from 2013 – 2017 within Papum RF. While 81% of the RF was under forest in 2013, it declined to around 76% in 4 years. The area under forest, as of winter 2017, is 561 km^2^ (Supplementary Figure 2). From 2013 to 2017, there was a loss of 32 km^2^ of forest, with an increase in logged-forest (27.22 km^2^) and of area under non-forest (4.76 km^2^). Out of a total area of 737 km^2^ classified, 156 km^2^ was logged-forest by 2017.

**Table 1.** Forest loss in the Papum Reserved Forest, Khellong Forest Division, Arunachal Pradesh quantified using RapidEye data for 2013, 2014, 2017. The total area of the Papum RF that was classified was 737 km^2^. Numbers in parentheses indicate percentages.

| Year | Logged-forest in km<sup>2</sup> (%) | Forest in km<sup>2</sup> (%) | Non-forest in km<sup>2</sup> (%) |
| --- | --- | --- | --- |
| 2013 | 128.8 (17.5) | 593.8 (80.8) | 14.3 (1.9) |
| 2014 | 166 (22.5) | 556.5 (75.5) | 14.3 (1.9) |
| 2017 | 156 (21.2) | 561 (76.2) | 19 (2.6) |

Our analyses recorded forest loss to be lower in 2017 than in 2014, for two reasons: (1) an area (~5 km^2^) in the eastern part of Papum RF was logged in 2014 but shows growth of secondary vegetation in 2017. The spectral nature of this 5 km^2^ area is very similar to forest and in 2017, the area is classified as forest. (2) Images in 2017 had a higher illumination elevation angle (46.05°), than in 2014 (39.48°), illuminating mountain slopes and forests that were previously under shadows. The illumination of river beds in 2017 also explains the increase in non-forest areas. The annual rate of forest area loss was 1.4 % year-1 corresponding to 8.2 km^2^ year-1.

### Forest loss around hornbill nests: 2011 – 2019

Forest area consistently dropped from 2011 to 2016, then increased in 2017, and decreased again up to 2019 (Fig. 2a). However, by 2019, only 45% of the 48 km^2^ of the 1-km buffer area around 29 hornbill nests was forested as compared to 80% in 2011 (Table 2). Forest loss is also evident from the construction of roads, burn scars and clear-cut felling of primary forest areas (Supplementary Figure 3). During the period from 2011 to 2015, the total forest loss around nest trees was about 6 km^2^, however this increased to a loss of 4 km^2^ in just one year in 2016, followed by a gain shown in 2017, with a loss of 8.59 km^2^ showing up in 2018 (Table 2). In the last 9 years, there has been a total loss of 16.61 km^2^ in a 1 km buffer around the 29 nest sites (Fig. 2b). Annual rate of forest area loss around the nest trees was 7% year-1, corresponding to 2.07 km^2^ year-1.

**Fig. 2.**
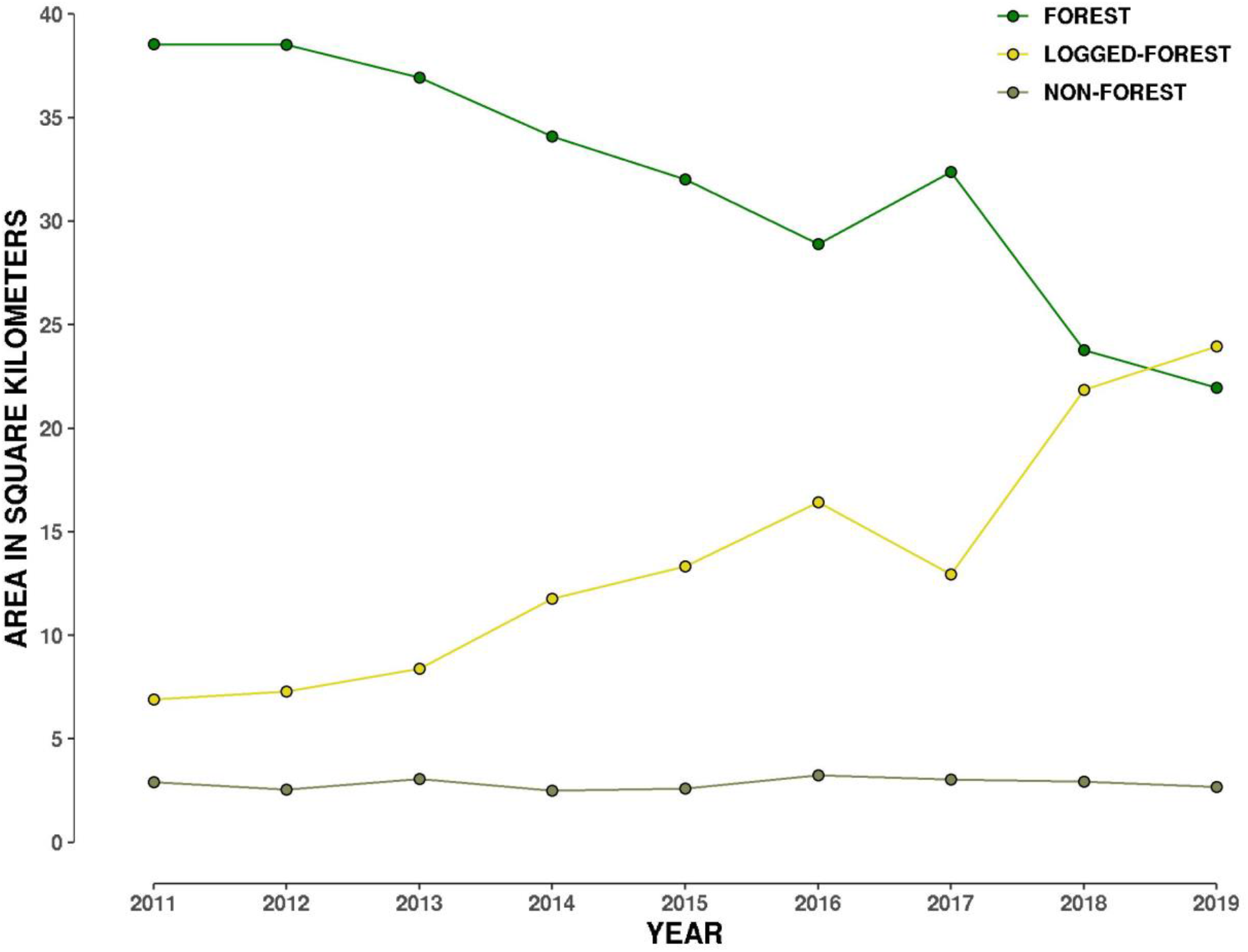
Plot showing change in area of three land-cover types from 2011 to 2019 within a 1 km buffer around the 29 hornbill nest sites.

**Table 2.** Forest loss around 29 hornbill nests in the Papum Reserved Forest, Khellong Forest Division, Arunachal Pradesh, north-east India; 2011 – 2019.

| Year | Forest area km<sup>2</sup> | % Forest area |
| --- | --- | --- |
| 2011 | 38.55 | 79.71 |
| 2012 | 38.52 | 79.66 |
| 2013 | 36.92 | 76.35 |
| 2014 | 34.09 | 70.49 |
| 2015 | 32.01 | 66.75 |
| 2016 | 28.90 | 59.50 |
| 2017 | 32.37 | 66.93 |
| 2018 | 23.78 | 48.95 |
| 2019 | 21.94 | 45.17 |

## Disscussion

The forest loss has serious consequences for tropical biodiversity, as the destruction of suitable habitat threatens the survival of forest specialist species (Tracewski et al. 2016). Several prior studies in the area have documented the negative effects of logging on key faunal groups, vegetation structure and composition, food abundance and seed dispersal (Datta 1998; Datta & Goyal 2008; Sethi & Howe 2009; Velho et al. 2012; Naniwadekar et al. 2015).

Selective logging on a commercial scale occurred in these Reserved Forests till the Supreme Court ban in 1996 (Datta 1998; Datta & Goyal 2008). Some level of illegal timber felling continued to occur in some pockets, however officially timber extraction for commercial purposes has been banned since 1996. Forest loss and degradation continued due to various other factors. Several current settlements existed prior to the declaration of the Reserved Forest, however population has grown subsequently leading to ambiguity and conflict in terms of people’s land rights and legal status of forests in the area. After devastating floods in May 2004, many families lost agricultural land to erosion, and some areas along the Assam-Arunachal border were occupied in anticipation of future needs. Over the last decade, most households stopped cultivating due to loss of land to floods and crop damage by elephants (Tewari et al 2017). Rubber and tea plantations also came up in the lower areas bordering Assam after 2007. These factors have led to some forest cover loss along the border areas in the 2001-2009 period. Apart from the forest loss due to these factors, till 2011-12, timber extraction in the Seijosa area was mainly for household needs and subsistence use by people.

Field observations/media reports show that tree felling increased after 2015 and coincided with the use of mechanized chainsaws and hired labour from Assam who camped in the forest. Reports of trucks transporting timber in the night and the use of various routes for covert transport of timber became more frequent after 2015. From 2017, there was construction of several link roads in the area and the clearing of tree cover near Jolly-Galoso area for the development of an herbal garden by Patanjali Ayurveda Limited in the area which also resulted in the forest loss. Since 2018, after road construction began, there is also loss of forest cover along the stretch from Pakke Kessang-Saibung.

The loss of 32 km^2^ of forest over 4 years within Papum RF is a cause for concern also because the area receives heavy rainfall often resulting in floods and landslides. The depletion of tropical forests in Papum RF severely threatens the future subsistence needs of the local and regional population. Although we do not explicitly test for these effects of deforestation, it is expected that landslides will increase if forest cover is lost at such a rapid rate (Bradshaw et al. 2007; Kumar & Bhagavanulu 2007; Horton et al. 2017; Stanley & Kirschbaum 2017). Soils along river valleys are destabilized accelerating river erosion rates (Horton et al. 2017) and amplifying flood risk and severity (Bradshaw et al 2007). In mountainous regions, deforestation weakens slopes exacerbating rainfall-triggered landslides (Kumar & Bhagavanulu 2007; Stanley & Kirschbaum 2017), significantly altering river sedimentation and geomorphology (Latrubesse et al. 2009), leading to cascading natural hazards like landslide dams.

Deforestation alters local climate resulting in drier, warmer conditions and reduced agricultural productivity (Lawrence & Vandecar 2014) and decreased access to clean drinking water (Mapulangaa & Naito, 2019). Furthermore, with climate change rapidly altering weather patterns, securing forests for their ecosystem services will be a pragmatic goal for all privileged and underprivileged stake-holders as per several sustainable development goals laid out by the United Nations.

### Possible effects of illegal logging on hornbills

The loss of around 35% of the forest area around the hornbill nest trees from 2011 to 2019 is alarming. From *ca*. 38 km^2^ in 2011, the area under forest declined to 21.94 km^2^ in 2019.

The forest cover change analysis shows that there has been loss and degradation of the surrounding nesting habitat and hornbill food trees. Although, the HNAP has protected individual hornbill nest trees and the immediate habitat surrounding the nest tree (Rane & Datta 2015) with an estimated 119 hornbill chicks have fledged from the protected nest trees from 2012 to 2018 (Parashuram & Datta 2018), the area within 1 km radius of nest trees has been considerably degraded by illegal logging.

This will likely have negative consequences for hornbill nesting and persistence in the Papum RF. Tree density/basal area and food and nest tree density is considerably lower in the RF than in the Pakke TR (Datta et al. unpublished data). An earlier study has documented the negative effects of logging on hornbills and vegetation structure and composition in the area (Datta 1998, Datta & Goyal 2008). Logging also reduces food abundances for hornbills and together with hunting has consequences for seed dispersal by hornbills (Sethi & Howe 2010, Naniwadekar et al. 2015b). In any case, while most of the earlier studies have all looked at the effects of ‘selective’ logging after some years since logging or when the logging was officially permitted before 1996, this study notes the alarming loss of forest despite the 1996 Supreme Court ban and the lack of any working plan under which the current logging is occurring within Papum RF.

Hornbills are highly mobile species with large home ranges, and nesting males move from the RF to the Pakke TR to forage for fruits. Our telemetry data of tagged Great and Wreathed hornbills show that some individual hornbills move between the Pakke TR and the RF (Naniwadekar et al. 2019b). However, despite their ability to move between these areas, a continuing loss of forest cover will result in nest trees in the RF becoming inactive. As the forest is becoming more degraded and is being logged it has also become more common to find only nests of the more adaptable Oriental Pied hornbill in the RF (Parashuram & Datta 2018), which is more common in open secondary forests (Datta 1998).

The tree felling occurs mainly in the drier months starting from September to March-April, but in some years, illegal logging activity has continued in the wetter period. March is the beginning of the breeding season for the larger-sized Great Hornbill and Wreathed hornbill when the females start entering the nest cavities, sealing them and laying eggs. Apart from the direct loss of forest habitat and individual trees, the sound of mechanized chainsaws, movement and presence of hired labour in camps and trucks results in disturbance during this critical time in the hornbill breeding season. It is likely that hornbill breeding is being negatively affected by the ongoing illegal logging activities which has increased in intensity in the last 2-3 years. Our long-term monitoring of hornbill roost sites located along the southern boundary of Pakke TR near the Pakke River, also shows movement of hornbills from Pakke TR to the Papum RF. The disturbance from illegal logging and loss of habitat, may also affect the use of roost sites by hornbills in the future.

## Conclusion

One of the challenges in our study was the strict classification of land-cover as non-forest and logged-forest. Our ROI includes areas that often flood in the monsoon changing the percentages of land-cover every year. New road construction or mining in recently logged forests can be classified as non-forest, while previously cleared primary forest can show regrowth as secondary vegetation. The difficult terrain in the region makes robust collection of ground-control points challenging. Hence, we make the following suggestions: 1) dry summer season images are best to distinguish secondary and non-woody vegetation from primary forest, 2) a binary classification system of forest and non-forest, and 3) forest loss estimations within a completely forested region such that loss in later years can be detected using year-to-year image subtraction techniques. However, we hope our work is a step towards achieving accurate forest loss estimates for an under-explored, mountainous region with exceptional forests and biodiversity.

The spurt in illegal commercial logging activities on a large-scale, with timber being sold and transported out of the state, using mechanized chainsaws and hired labourers from a neighbouring community, is driving an alarming loss of forest cover in this area. In addition, with the construction of new roads, the continuation of these illegal activities to newer areas in the higher northern parts of the RF deeper inside Arunachal Pradesh is also being facilitated and is a threat to the long-term status of this important forest area for both people and wildlife.

We recommend the following management measures to stop the illegal logging are 1) a complete ban on the use/sale and possession of mechanized chainsaws in the area. While prohibitory orders have been issued in the past by the district administration, these have not been enforced, 2) stopping the unregulated movement of hired labour from the neighbouring state into Arunachal Pradesh for their use in illegal logging and transportation activities, 3) a thorough on-ground survey of the areas affected along with official and transparent records of seizure and disposal of seized timber from inside the forest and from illegal timber depots 3) night patrolling by police/Forest Department staff on all possible movement routes to stop the movement of trucks carrying timber out of the state, 4) the establishment of regular forest and/or community monitoring patrols to check illegal felling within the RF and 5) a constant monitoring of the state of forest cover by an external agency to ensure that illegal logging has been stopped. In the long-term, for better governance, clarity in the use and ownership of forest land also needs to be addressed under the law given that some of the designated forest area is under settlements and multiple use areas by people.

## Code Availability

The code for image classification is publicly available on https://github.com/monsoonforest/deforestation/blob/master/randomForest-image-classification.

## Data Availability

RapidEye and PlanetScope datasets are not openly available as Planet Labs is a commercial company. CS obtained the datasets through Planet Lab’s Education and Research program upon application. The classified land-cover datasets can be made available upon request from the authors.

## Acknowledgments

We thank Rohit Naniwadekar, TR Shankar Raman, Divya Mudappa, Kulbhushuan Suryawanshi, Charudutt Mishra for comments on earlier drafts of the paper. We thank the field staff and nest protectors from the Nyishi community for monitoring and protecting the hornbill nests. CS is grateful to M. Raghurama, S. Virdi and S. Pulla for suggestions that improved the analyses. We are grateful to Planet Labs for providing free access to their data to CS via their education and research program. We are grateful to J-P Puyravaud for reviewing the manuscript and for his valuable comments.

## Author contributions

AD and CS conceived the idea and the study; DP and AD provided field data; CS analysed the data; AD and CS wrote the paper with inputs from DP.

## References

Ahmad W, Tahseen Q, Baniyamuddin M, Hussain A (2004) Description of two new species of Plectinae (Nematoda: Araeolaimida) from India. Nematology 6:755–764.

Ahti T, Dixit PK, Singh KP, Sinha GP (2002) *Cladonia singhii* and other new reports of *Cladonia* from the Eastern Himalayan Region of India. The Lichenologist 34:305–310.

Athreya R (2006) A new species of Liocichla (Aves: Timallidae) from Eaglenest Wildlife Sanctuary, Arunachal Pradesh, India. Indian Birds 2: 82–94.

Anonymous (2017) Illegal logging on the rise in Arunachal PTI, India Today, November 24, 2017. https://www.indiatoday.in/pti-feed/story/illega1-1ogging-on-the-rise-in-amnacha1-1093159-2017-11-24.

Anonymous (2019) Illegal wooden logs worth Rs 4 lakh seized from LPG carrying truck in Tezpur. The Sentinel, Assam. 28 January, 2019. https://www.sentinelassam.com/news/illega1-wooden-1ogs-worth-rs-4-1akh-seized-from-1pg-carrying-truck-in-tezpur

Bradshaw CJA, Sodhi NS, Peh KSH, Brook BW (2007) Global evidence that deforestation amplifies flood risk and severity in the developing world. Global Change Biology 13:2379–2395.

Census of India (2011) States Census 2011. http://censusindia.gov.in/ [Accessed 15 June 2019].

Captain A, Deepak V, Pandit R, Bhatt B, Athreya R. 2019. A new species of pitviper (Serpentes: Viperidae: *Trimeresurus* Lacepède, 1804) from West Kameng district, Arunachal Pradesh, India. Russian Journal of Herpetology 26:111–122.

Curtis PG, Slay CM, Harris NL, Tyukavina A, Hansen MC (2018) Classifying drivers of global forest loss. Science 361:1108–1111. https://doi.org/10.1126/science.aau3445.

Dalvi S (2013) Elliot’s Laughing thrush *Trochalopteron elliotii* and Black-headed Greenfinch *Chloris ambigua* from Anini, Arunachal Pradesh, India. Indian Birds 8:130.

Dasgupta S, Hilaluddin (2012) Differential effects of hunting on populations of hornbills and imperial pigeons in the rainforests of the Eastern Indian Himalaya. Indian Forester 138:902–909.

Datta A (1998) Hornbill abundance in unlogged forest, selectively logged forest and a plantation in western Arunachal Pradesh. Oryx 32:285–294.

Datta A (2001) An ecological study of sympatric hornbills and fruiting patterns in a tropical forest in Arunachal Pradesh. 245 pp. Ph.D Thesis submitted to Saurashtra University, Rajkot, Gujarat (affiliate of Wildlife Institute of India).

Datta A, Rawat GS (2003) Foraging patterns of sympatric hornbills in the non-breeding season in Arunachal Pradesh, north-east India. Biotropica 35:208–218.

Datta A, Rawat GS (2004) Nest site selection and nesting success of hornbills in Arunachal Pradesh, north-east India. Bird Conservation International 14:249–262.

Datta A, Goyal SP (2008) Responses of diurnal squirrels to selective logging in western Arunachal Pradesh. Current Science 95:895–902.

Datta A, Rane A, Tapi T (2012) Shared parenting: Hornbill Nest Adoption Program in Arunachal Pradesh. The Hindu Survey of the Environment pp. 88–97.

Datta A, Naniwadekar R (2015) Hope for hornbills. In: Hegan A (ed) No more endlings: Saving species one story at a time. Coalition Wild and The Wild Foundation.

Forest Survey of India (2005) State of Forest Report 2005. Ministry of Environment and Forests, Government of India, Dehra Dun, India.

Forest Survey of India (2009) State of Forest Report 2009. Ministry of Environment and Forests, Government of India, Dehra Dun, India.

Forest Survey of India (2011) State of Forest Report 2011. Ministry of Environment and Forests, Government of India, Dehra Dun, India.

Forest Survey of India (2013) State of Forest Report 2013. Ministry of Environment and Forests, Government of India, Dehra Dun, India.

Forest Survey of India (2015) State of Forest Report 2015. Ministry of Environment and Forests, Government of India, Dehra Dun, India.

Forest Survey of India (2017) State of Forest Report 2017. Ministry of Environment and Forests, Government of India, Dehra Dun, India.

Gajurel PR, Rethy P, Kumar Y (2001) A new species of Piper (Piperaceae) from Arunachal Pradesh, north-eastern India. Botanical Journal of the Linnean Society 137:417–419.

Gibbs HK, Ruesch AS, Achard F, Clayton MK, Holmgren P, Ramankutty N, Foley JA (2010) Tropical forests were the primary sources of new agricultural land in the 1980s and 1990s. Proceedings of the National Academy of Science USA 107:16732–16737.

Gibson L, Lee TM, Koh LP, Brook BW, Gardner TA, Barlow J, Peres CA, Hansen MC, Potapov PV, Moore R, Hancher M, Turubanova SA, Tyukavina A, Thau D, Stehman SV, Goetz SJ, Loveland TR, Kommareddy A, Egorov A, Chini L, Justice CO, Townshend, JRG (2013) High-resolution global maps of 21st-century forest cover change. Science 342:850–853. http://earthenginepartners.appspot.com/science-2013-global-forest/download_v1.6.html.

Global Forest Watch (2019) “Tree Cover Loss in India”. Accessed on 25th May 2019 from www.globalforestwatch.org.

Hareesh VS, Gogoi R, Sabu M (2016) *Impatiens pseudocitrina* (Balsaminaceae), a new species from Arunachal Pradesh, northeast India. Phytotaxa 282:231–234.

Horton AJ, Constantine JA, Hales TC, Goossens B, Bruford MW, Lazarus ED (2017) Modification of river meandering by tropical deforestation. Geology 45:511–514.

The Indian Forest Act (1927) Act XVI of 1927 (as modified up to 15 June 1951) Govt of India. http://extwprlegs1.fao.org/docs/pdf/ind3171.pdf [Accessed 17 June 2019]

International Union for Conservation of Nature (2018) The IUCN Red List of Threatened Species 2018.

Khandekar N. 2019. Between tradition and trafficking: opium in Arunachal. https://www.thethirdpole.net/en/2019/05/08/between-tradition-and-trafficking-opium-in-arunachal-pradesh/. May 8, 2019.

Kumar SV, Bhagavanulu DVS (2008) Effect of deforestation on landslides in Nilgiris district - A case study. Journal of the Indian Society of Remote Sensing 36:105.

Kushwaha SP, Hazarika R (2004) Assessment of habitat loss in Kameng and Sonitpur Elephant Reserves. Current Science 87:1447–1453.

Latrubesse EM, Amsler ML, de Morais RP, Aquino S (2009) The geomorphologic response of a large pristine alluvial river to tremendous deforestation in the South American tropics: The case of the Araguaia River. Geomorphology 113: 239–252.

Lawrence D, Vandecar K (2015) Effects of tropical deforestation on climate and agriculture. Nature Climate Change 5: 27.

Liaw, A, Wiener M (2002). Classification and Regression by random Forest. R News 2(3): 18–22.

Mahony S, Kamei RG, Teeling EC, Biju SD (2018) Cryptic diversity within the Megophrys major species group (Amphibia: Megophryidae) of the Asian Horned Frogs: Phylogenetic perspectives and a taxonomic revision of South Asian taxa, with descriptions of four new species. Zootaxa 4523:1–96.

Mamai J (2018) Rampant destruction of forests in Namdang. Arunachal Times, November 26, 2018. https://arunachaltimes.in/index.php/2018/11/26/rampant-destruction-of-forests-in-namdang/

Mazoomdar J (2011) Where the forests have no trees. http://www.openthemagazine.com/article/nation/where-the-forests-have-no-trees/. Accessed 19 May 2013.

Milodowski DT, Mitchard ETA, Williams M (2017) Forest loss maps from regional satellite monitoring systematically underestimate deforestation in two rapidly changing parts of the Amazon. Environmental Research Letters 12:094003.

Mittermeier RA, Myers M, Mittermeier, CG (2000) Hotspots: earth’s biologically richest and most endangered terrestrial ecosystems. Conservation International, Mexico.

Mishra C, Datta A (2007) A new bird species from Eastern Himalayan Arunachal Pradesh – India’s biological frontier. Current Science 92:1205–06.

Manly, B. F. (1991) Randomization, bootstrap and Monte Carlo methods in biology. Chapman and Hall/CRC.

Mapulanga AM, Naito H (2019) Effect of deforestation on access to clean drinking water. Proceedings of the National Academy of Sciences 116:8249–8254.

Naniwadekar R, Mishra C, Isvaran K, Madhusudan MD, Datta A (2015a) Looking beyond parks: the conservation value of unprotected area for hornbills in Arunachal Pradesh, Eastern Himalaya. Oryx 49:303–311.

Naniwadekar R, Shukla U, Isvaran K, Datta A (2015b) Reduced hornbill abundance associated with low seed arrival and altered recruitment in a hunted and logged tropical forest. PLoS ONE DOI: 10.371/journal.pone.0120062.

Naniwadekar R, Chaplod S, Datta A, Rathore A, Sridhar, H (2019a) Large frugivores matter: insights from network and seed dispersal effectiveness approaches. Journal of Animal Ecology DOI: 10.1111/1365-2656.13005.

Naniwadekar R, Rathore, A, Shukla, U, Chaplod, S, Datta A (2019b) How far do Asian hornbills disperse seeds? Acta Oecologica 101(2019) 103482, https://doi.org/10.1016/j.actao.2019.103482.

National Tiger Conservation Authority of India (2012) https://projecttiger.nic.in/content/109_1_ListofTigerReservesCoreBufferAreas.aspxref) [accessed 15 June 2019].

Neary DG, Ryan KC, DeBano LF, eds. 2005. (revised 2008). Wildland fire in ecosystems: effects of fire on soils and water. Gen. Tech. Rep. RMRS-GTR-42-vol.4. Ogden, UT: U.S. Department of Agriculture, Forest Service, Rocky Mountain Research Station. 250 p.

Olofsson P, Foody GM, Herold M, Stehman SV, Woodcock CE, Wulder MA (2014) Good practices for estimating area and assessing accuracy of land change. Remote Sens. Environ. 148: 42–57.

Pandit, MK, Sodhi NS, Koh LP, Bhaskar A, Brook BW (2007) Unreported yet massive deforestation driving loss of endemic biodiversity in Indian Himalaya. Biodiversity and Conservation 16:153–163.

Parashuram D, Datta A (2018) Hornbill Nest Adoption Program Report. 13 pp. http://ncf-india.org/projects/hornbill-nest-adoption-program.

Planet Labs Incorporate (2019) Planet imagery product specifications. August 2019. https://assets.planet.com/docs/combined-imagery-product-spec-final-august-2019.pdf

Praveen J, Jayapal R, Pittie A (2016) A checklist of the birds of India. Indian Birds 11:113–172.

Praveen J, Jayapal R, Pittie A (2019) Checklist of the birds of India (v2.3). Website: http://www.indianbirds.in/india/ [Date of publication: 15 January, 2019].

Purkayastha J, David P. 2019. A new species of the snake genus *Hebius* Thompson from Northeast India (Squamata: Natricidae). Zootaxa 4555:79–90.

Puyravaud JP (2003) Standardizing the calculation of the annual rate of deforestation. Forest Ecology and Management, 177: 593–596.

Puyravaud JP, Davidar P, Laurance WF (2010) Cryptic destruction of India’s native forests. Conservation Letters 3:390–394.

Puyravaud JP, Davidar P, Laurance WF (2010) Cryptic loss of India’s native forests. Science 329:32.

QGIS Development Team (2019) QGIS Geographic Information System. Open Source Geospatial Foundation Project. http://qgis.osgeo.org.

R Core Team (2016) R: A language and environment for statistical computing. R Foundation for Statistical Computing, Vienna, Austria. https://www.R-project.org/.

Rane A, Datta A (2015) Protecting a hornbill haven: a community-based conservation initiative in Arunachal Pradesh, north-east India. Malayan Nature Journal 67:203–18.

Rina T (2017) Large-scale timber logging in Papum Reserved Forest. Arunachal Times, April 20, 2017. https://www.arunachaltimes.in/archives/apr17_20.html.

Rina T (2019) NGT steps in on illegal logging in Papum Reserved Forest. Arunachal Times, April 17, 2019. https://arunachaltimes.in/index.php/2019/04/17/ngt-steps-in-on-illegal-logging-in-papum-reserve-forest/

Roy P (2013) *Callerebia dibangensis* (Lepidoptera: Nymphalidae: Satyrinae), a new butterfly species from the eastern Himalaya, India. Journal of Threatened Taxa 5:4725–4733.

Sethi P, Howe HF (2009) Recruitment of hornbill-dispersed trees in hunted and logged forests of the Indian Eastern Himalaya. Conservation Biology 23:710–718.

Sekhar S (2014a) Disappearing oasis: north-eastern India losing forests as people move in. 18 November 2014. Mongabay.com https://news.mongabay.com/2014/11/disappearing-oasis-northeastern-india-losing-forests-as-people-move-in/

Sekhar S (2014b) Conflict-fueled deforestation, poaching in Assam continue despite truce. 19 November 2014. Mongabay.com https://news.mongabay.com/2014/11/conflict-fueled-deforestation-poaching-in-assam-continue-despite-truce/

Siliwal M, Molur S, Raven R (2015) New genus with two new species of the family Nemesiidae (Araneae: Mygalomorphae) from Arunachal Pradesh, India. Journal of Asia-Pacific Biodiversity 8:43–48.

Sinha A, Datta A, Madhusudan MD, Mishra C (2005) *Macaca munzala:* a new species from western Arunachal Pradesh, northeastern India. International Journal of Primatology 26:977–989.

Sodhi NS, Koh LP, Brook BW, Ng PKL (2004) Southeast Asian biodiversity: an impending disaster. Trends in Ecology and Evolution 19:654–660.

Sondhi S, Ohler A (2011) A blue-eyed *Leptobrachium* (Anura: Megophryidae) from Arunachal Pradesh, India. Zootaxa 2912:28–36.

Srinivas A (2018) India’s forest cover: What data shows. Live Mint 4 July 2018 https://www.livemint.com/Politics/jUW0iY07OS0mRMi1gI5YzH/India-forest-cover-What-data-shows.html. Accessed 15 April, 2019.

Srinivasan U (2014) Oil Palm Expansion: Ecological threat to north-east India. Economic and Political Weekly 49(36) Sep 8, 2014.

Srinivasan U (2018) Marginalisation, migration and militancy: the complexities of forest and biodiversity loss on the Assam-Arunachal border. In: Srinivasan U & Velho N (eds) Conservation from the Margins. Orient Black Swan. Hyderabad, pp 177–197.

Srivastava S, Singh TP, Singh H, Kushwaha SPS, Roy PS (2002) Assessment of large-scale deforestation in Sonitpur district of Assam. Current Science 82:1479–1484

Stanley T, Kirschbaum DB (2017) A heuristic approach to global landslide susceptibility mapping. Natural Hazards 87: 145–164

Tamang L, Chaudhry S, Choudhury D (2008) *Erethistoides senkhiensis*, a new catfish (Teleostei: Erethistidae) from India. Ichthyological Exploration of Freshwaters 19:185–191

Teegalapalli K, Datta A (2016) Shifting to settled cultivation: changing practices among the *Adis* in Central Arunachal Pradesh, north-east India. Ambio 45:602–612 https://doi.org/10.1007/s13280-016-0765-x.

Teegalapalli K, Datta A (2017) Top-down or bottom-up: the role of government and local institutions in regulating shifting cultivation in the Upper Siang district, Eastern Himalaya, India. Pages 760–766 In: Shifting Cultivation Policies: Balancing environmental and social sustainability, Edited by Cairns, M., Routledge, UK.

Tiwari SK, Kyarong S, Choudhury C, Williams AC, Ramkumar K, Deori D (2017) Elephant Corridors of North-Eastern India. Pages 424–573 In: Right of Passage: Elephant Corridors of India (2nd Edition). Menon V, Tiwari SK, Ramkumar K, Kyarong S, Ganguly U, Sukumar, R (eds). Conservation Reference Series No. 3. Wildlife Trust of India, New Delhi.

Tucker CJ, Grant DM, Dykstra JD (2004). NASA’s global ortho-rectified Landsat data set. Photogrammetric Engineering & Remote Sensing, 70: 313–322.

Tracewski L, Butchart SHM, Marco MD, Ficetola GF, Rondinini C, Symes A, Wheatley H, Beresford, AE, Buchanan GM (2016) Towards quantification of the impact of 21st century deforestation on the extinction risk of terrestrial vertebrates. Conservation Biology 30:1070–1079.

Velho N, Srinivasan U, Prashanth NS, Laurance WF (2011) Human disease hinders anti-poaching efforts in Indian nature reserves. Biological Conservation 144:2382–2385. https://doi.org/10.1016/j.biocon.2011.06.003

Velho N, Agarwala M, Srinivasan U, Laurance WF (2014) Collateral damage: impacts of ethno-civil strife on biodiversity and natural resource use near Indian nature reserves. Biodiversity and Conservation 23:2515–2527.

Velho N, Datta A, Datta-Roy A (2016) An inclusive oil palm policy for people and biodiversity. The Arunachal Times, November 9, 2016. http://www.arunachaltimes.in/an-inclusive-oil-palm-policy-for-people-and-biodiversity/.

Wasteland Atlas 2011. Wastelands Atlas of India, prepared by National Remote Sensing Centre, Department of Land Resources, Ministry of Rural Development, India.

Zanan RL, Nadaf AB (2012) *Pandanus martinianus* (Pandanaceae), a new endemic species from northeastern India. Phytotaxa 73:1–7.

