## Supplementary tables and figures for "Persistent loss of biologically-rich tropical forests in the Indian Eastern Himalaya" (from another version of this preprint: version 2, 2019-11-12)

^1^Chintan Sheth

National Centre for Biological Sciences

Tata Institute of Fundamental Research

Bellary Road

Bangalore 560065

Karnataka, India

^2^Aparajita Datta

Nature Conservation Foundation

1311,“Amritha”, 12th Main
Vijayanagar 1st Stage
Mysore 570 017

Karnataka, India

^2^Devathi Parashuram

Nature Conservation Foundation

1311, “Amritha”, 12th Main
Vijayanagar 1st Stage
Mysore 570 017

Karnataka, India

**Corresponding author**

**Aparajita Datta**

Ph: +91 9448351560

**Co-corresponding author**

**Chintan Sheth**

Ph: +91 9449056489

**Supplementary Table 1:** Details of forest cover and change (in square kilometers) from 2003 to 2017 in Arunachal Pradesh state, north-east India. Data from FSI reports 2003-2017.

| Assessment year | Very dense forest | Moderately dense forest | Open forest | Total forest | Percentage of area | Net change | Net actual change |
| --- | --- | --- | --- | --- | --- | --- | --- |
| 2003 | 14,445 | 38,084 | 15,163 | 67,692 | 80.83 | -26 | -26 |
| 2005 | 14,411 | 37,977 | 15,389 | 67,777 | 80.93 | 85 | 85 |
| 2007 | ND | ND | ND | ND | ND | ND | ND |
| 2009 | 20,858 | 31,556 | 14,939 | 67,353 | 80.43 | -424 | -119* |
| 2011 | 20,868 | 31,519 | 15,023 | 67,410 | 80.5 | 57 | -74* |
| 2013 | 20,828 | 31,414 | 15,079 | 67,321 | 80.39 | -89 | -89 |
| 2015 | 20,806 | 31,181 | 15,167 | 67,154 | 80.3 | -167 | -73* |
| 2017 | 20,721 | 30,955 | 15,288 | 66,964 | 79.96 | -190 | -190 |

* Modifications and corrections based on re-interpretation and checking of satellite data, as mentioned in the cited FSI reports (2003-2017). Assessment year is the year of the FSI report, but for some years, the data may be from previous years.

**Supplementary Table 2:** Tree cover loss within Arunachal Pradesh from 2001-2018. Data from Global Forest Watch (2019).

| **Year** | **Loss of > 30% tree cover (km^2^)** |
| --- | --- |
| 2001 | 86.03 |
| 2002 | 70.1 |
| 2003 | 64.09 |
| 2004 | 128.04 |
| 2005 | 88.68 |
| 2006 | 98.02 |
| 2007 | 95.87 |
| 2008 | 110.15 |
| 2009 | 90.85 |
| 2010 | 59.37 |
| 2011 | 124.64 |
| 2012 | 143.66 |
| 2013 | 92.81 |
| 2014 | 130.5 |
| 2015 | 123.65 |
| 2016 | 190.16 |
| 2017 | 190.2 |
| 2018 | 113.52 |
| **Total** | **2000.34** |

**Supplementary Figure 1:** Forest cover change from the period of 2003-2011. Data from FSI reports 2003-2017. There was no report/data for 2007.

Forest cover

Area in sq. km

|  | Forest cover |
| --- | --- |
| 2003 | 67692 |
| 2005 | 67777 |
| 2009 | 67353 |
| 2011 | 67410 |
| 2013 | 67321 |
| 2015 | 67154 |
| 2017 | 66964 |

**Supplementary Table 3:** List of images, year and the spatial extent of the classification of the image.

| **IMAGE NUMBER** | **DATASET** | **YEAR** | **EXTENT OF SPATIAL ANALYSIS** |
| --- | --- | --- | --- |
| 4651515_2011-01-07_RE3_3A_Analytic | RapidEye | 2011 | Nest Sites within Papum RF |
| 4651515_2012-12-25_RE1_3A_Analytic | RapidEye | 2012 | Nest Sites within Papum RF |
| 4651514_2013-11-14_RE2_3A_Analytic | RapidEye | 2013 | Papum RF |
| 4651515_2013-11-14_RE2_3A_Analytic | RapidEye | 2013 | Nest sites and full extent of Papum RF |
| 4651516_2013-11-14_RE2_3A_Analytic | RapidEye | 2013 | Papum RF |
| 4651614_2013-11-14_RE2_3A_Analytic | RapidEye | 2013 | Papum RF |
| 4651615_2013-11-14_RE2_3A_Analytic | RapidEye | 2013 | Papum RF |
| 4651514_2014-12-29_RE2_3A_Analytic | RapidEye | 2013 | Papum RF |
| 4651515_2014-12-29_RE2_3A_Analytic | RapidEye | 2014 | Nest sites and full extent of Papum RF |
| 4651516_2014-12-29_RE2_3A_Analytic | RapidEye | 2014 | Papum RF |
| 4651614_2014-12-29_RE2_3A_Analytic | RapidEye | 2014 | Papum RF |
| 4651615_2014-12-29_RE2_3A_Analytic | RapidEye | 2014 | Papum RF |
| 4651515_2015-11-21_RE1_3A_Analytic | RapidEye | 2015 | Nest Sites within Papum RF |
| 20161212_061349_0c46_3B_AnalyticMS | PlanetScope | 2016 | Nest Sites within Papum RF |
| 20161212_061350_0c46_3B_AnalyticMS | PlanetScope | 2016 | Nest Sites within Papum RF |
| 20161212_061351_0c46_3B_AnalyticMS | PlanetScope | 2016 | Nest Sites within Papum RF |
| 4651514_2017-11-04_RE2_3A_Analytic | RapidEye | 2017 | Papum RF |
| 4651515_2017-11-04_RE2_3A_Analytic | RapidEye | 2017 | Nest sites and full extent of Papum RF |
| 4651516_2017-11-04_RE2_3A_Analytic | RapidEye | 2017 | Papum RF |
| 4651614_2017-11-04_RE2_3A_Analytic | RapidEye | 2017 | Papum RF |
| 4651615_2017-11-04_RE2_3A_Analytic | RapidEye | 2017 | Papum RF |
| 20180512_035145_100a_3B_AnalyticMS_SR | PlanetScope | 2018 | Nest Sites within Papum RF |
| 20180512_035146_100a_3B_AnalyticMS_SR | PlanetScope | 2018 | Nest Sites within Papum RF |
| 20190425_032704_0f36_3B_AnalyticMS_SR | PlanetScope | 2019 | Nest Sites within Papum RF |
| 20190425_032705_0f36_3B_AnalyticMS_SR | PlanetScope | 2019 | Nest Sites within Papum RF |
| 20190425_032706_0f36_3B_AnalyticMS_SR | PlanetScope | 2019 | Nest Sites within Papum RF |

**Supplementary Table 4:** Summary statistics (mean and standard deviation) for RapidEye and PlanetScope based forest loss maps after a manual inspection of points retrieved from a stratified random sample. The procedure was repeated for the images covering the full extent of Papum RF as well as images only used for the prediction within nest sites. NF = stable non-forest, F = stable forest and LF = logged-forest during the given time period. User's Accuracy is the proportion of the area mapped as class *i* that has a reference class *i*. Producer's Accuracy is the proportion of the area of reference class *i* that is mapped as class *i*.

| **SPATIAL SCALE OF ANALYSIS** | **Papum RF**  **2013 - 2017** | **Nest Sites within Papum RF**  **2011-2019** |
| --- | --- | --- |
| Number of points sampled per reference class | 700 | 700 |

| Overall Accuracy | 98.3 ± 0.8 % | 95.2 ± 1.3 % |
| --- | --- | --- |

|  | NF | F | LF | NF | F | LF |
| --- | --- | --- | --- | --- | --- | --- |
| User’s Accuracy | 96.8 ± 0.8 % | 99.9 % | 78.4 ± 3.8 % | 94.7± 3.3 % | 96.9 ± 1.8 % | 79.9 ± 3.2 % |
| Producer’s Accuracy | 94.7 ± 1.4 % | 99.0 ± 0.1% | 93.7 ± 1.8 % | 95.8 ± 1.2 % | 96 ± 1.3 % | 93.2 ± 1.8 % |

**Supplementary Table 5:** Confusion matrices for classified maps derived from a stratified random sample of test points for Papum RF and nest-sites with Papum RF, indicating the accuracy with which changing forest extent and deforestation is mapped for the periods 2013–2017 and 2011-2019 respectively.

| **Papum RF**  **2013-2017** | | Reference | | | Mapped Area Fraction |
| --- | --- | --- | --- | --- | --- |
|  |  | Logged Forest | Forest | Non-Forest |  |
| Map | Logged Forest | 0.044 | 0.009 | 0.004 | 0.056 |
|  | Forest | 0.001 | 0.876 | 0.000 | 0.877 |
|  | Non-Forest | 0.002 | 0.000 | 0.065 | 0.067 |
| Reference Area Fraction | | 0.047 | 0.885 | 0.068 | 1.000 |
| **Nest Sites within Papum RF**  **2011-2019** | | Reference | | | Mapped Area Fraction |
|  |  | CF | F | NF |  |
| Map | Logged Forest | 0.141 | 0.028 | 0.007 | 0.176 |
|  | Forest | 0.016 | 0.634 | 0.000 | 0.650 |
|  | Non-Forest | 0.003 | 0.005 | 0.166 | 0.174 |
| Reference Area Fraction | | 0.160 | 0.667 | 0.174 | 1.000 |

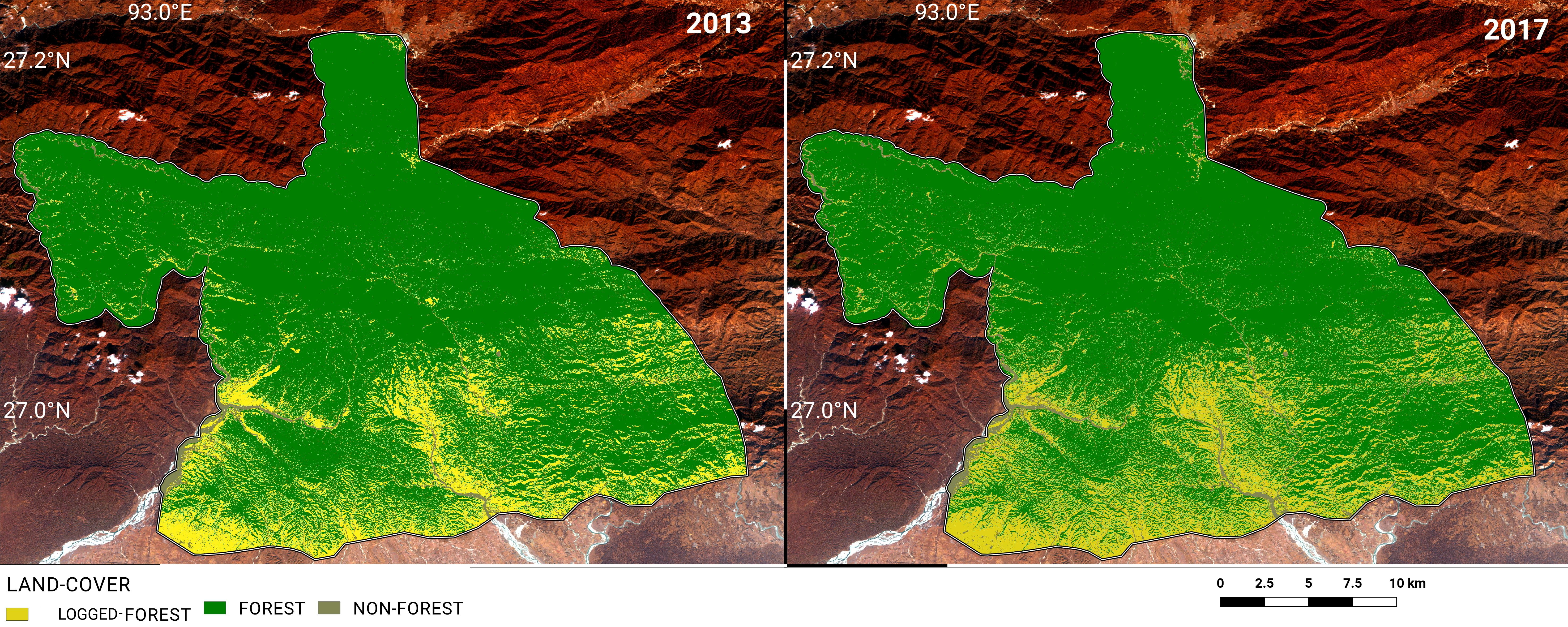

**Supplementary Figure 2**: Comparison of classified images of Papum RF (plus a 500m buffer around the boundary). The ‘salt and pepper’ effect of deforestation, is evident in the southern regions especially in 2017. A colour coded classified image is overlaid on a RapidEye false colour composite image using the Red Edge (B4; 690 – 730 nm), Green (B2; 520 – 590 nm) and Blue (B1; 440 – 510 nm) spectral bands. Colours in the spectral image are the same as described in the caption of figure 1 in main text.

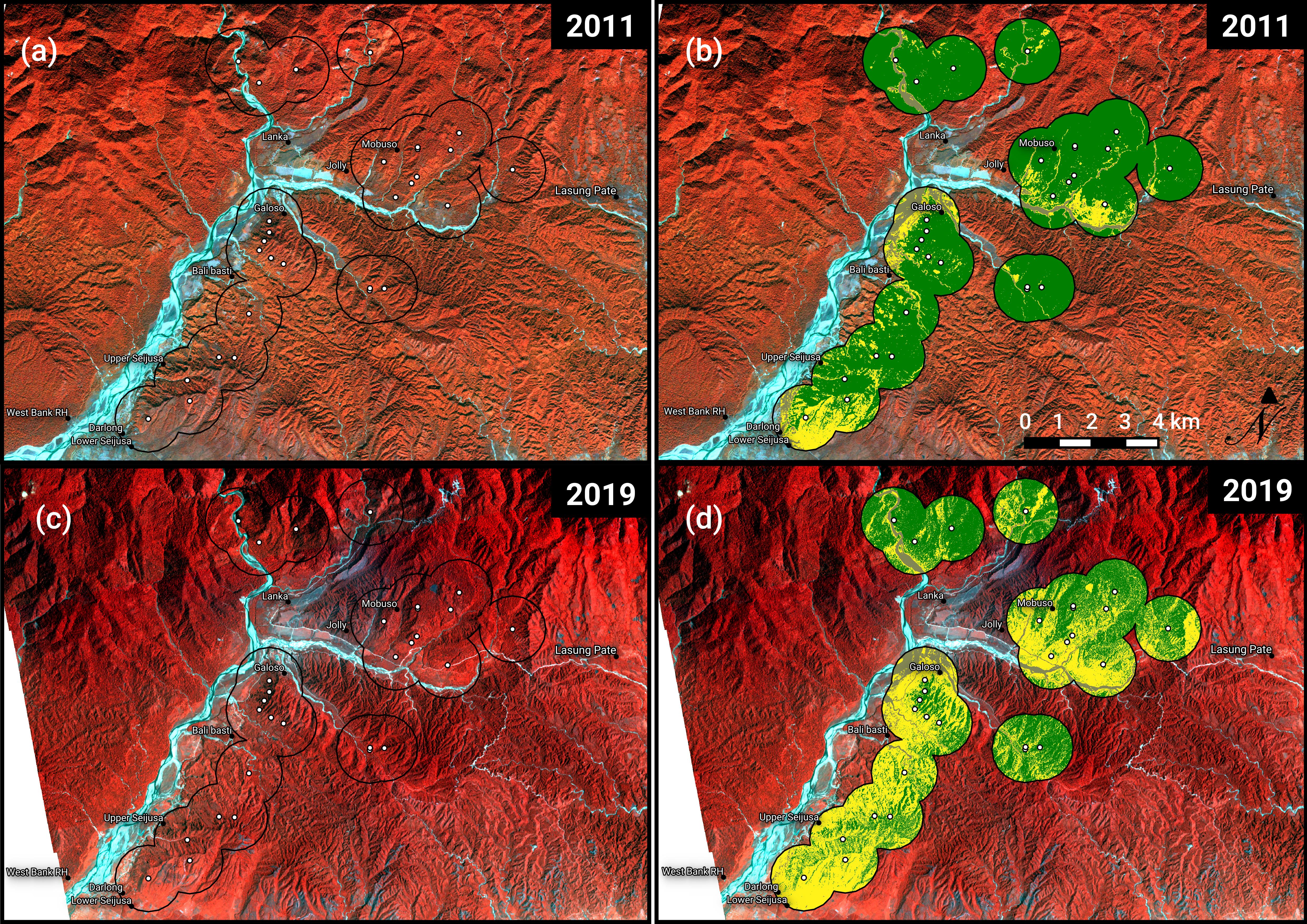

**Supplementary Figure 3:** Comparison of images between 2011 (a & b) and 2019 (c & d) of the nest sites. Images a. and c. are false colour composites of RapidEye (2011; RGB composite: band 4, band 2, band 1) and PlanetScope (2019; RGB composite: band 4, band 2, band 3) images. Panels b. and d. are the same false colour composites of the respective year, with an overlay of the classified image of the nest sites. Image classification colour key: yellow: logged-forest, green: forest and grey: non-forest. White dots enclosed within black circles are nest sites monitored by the Hornbill Nest Adoption Program. Black dots are villages. In the 2018 images, note the ash-grey areas on the southern hill slopes, indicative of fire scars. Surfaces with dark red indicated forest cover, lighter shades are either of cleared forest, plantations or secondary vegetation. Landslides and river beds are white. In the 2011 images, primary forest is dark orange to red. Secondary vegetation are bright shades of red while logged-forest areas are grey interspersed with red.
